# Mechanisms and Plasticity in Leaves and Leaflets in a Creeping Legume, *Mimosa pudica*

**DOI:** 10.1101/2025.05.03.652067

**Authors:** Mason T. Kellogg

**Author notes:** EAP Tropical Biology and Conservation Program, Spring 2024.

## Abstract

Mimosa pudica (Fabaceae) is a creeping plant known for its rapid thigmonastic movement upon touch, facilitated by specialized joint-like thickenings called pulvini. This study examines the activation behavior of the primary pulvinus (P1) in response to repeated touch stimuli, providing evidence for a mechanical exhaustion mechanism underlying the response. Experiments were conducted on M. pudica in Cuajiniquil, Costa Rica. Petiole angle change was recorded following repeated P1 stimuli, both with (P3 group) and without (NS-P3 group) concurrent tertiary pulvinus (P3) activation. Results showed the highest mean petiole angle change and P1 activations at the first stimulus, with a significant decline at the second stimulus and sustained lower responses thereafter. Both the NS-P3 and P3 groups exhibited similar overall behavior, characterized by a sharp decline in petiole angle change and P1 activation counts after the first stimulus. However, the P3 group had a lower initial petiole angle change compared to the NS-P3 group, and exhibited significant wave-like behavior, suggesting a more pronounced refractory period due to the combined activation of both P1 and P3 pulvini. The findings support a mechanical exhaustion explanation for the primary pulvinus behavior over repeated stimuli, where the rapid decline and sustained low responses suggest energy depletion and slow ion channel reset.

## INTRODUCTION

Plants experience herbivory which damages leaves and interferes with photosynthesis. Some plants may employ methods for defense against herbivores. One such example is thigmonastic movement wherein a plant will rapidly move in response to a touch stimulus. This action requires energetically costly active ion transport, but is theorized to deter herbivores (Fleurat-Lessard et al. 1997). Plants with this response maintain a balance between energy conservation and defense (Jensen 2011). *Mimosa pudica* is a creeping plant with pinnately compound leaves. Like many Fabaceae, it has pulvini – joint-like thickenings that facilitate growth-independent movement. *M. pudica* experiences a thigmonastic response to touch stimuli facilitated by its pulvini.

*M. pudica* features three types of pulvini (Figs 1 & 2). The primary pulvinus (P1), located at the base of each petiole, triggers the drooping of the entire compound leaf toward the stem base within approximately two to five seconds when activated. This P1 movement is believed to scare away potential herbivorous insects on the leaf (Minorsky 2019). Secondary pulvini (P2), situated at the base of each leaf’s rachis, are less sensitive to touch and less studied, yet they likely play a role in the closure of the pinnae. The tertiary pulvinus (P3), found at the base of each pinnule, causes the pinnule to fold up in response to touch within two to seven seconds. The adaptive significance of this P3 folding behavior is theorized to (1) scare away potential herbivorous insects on the leaf and (2) make the plant appear smaller and wilted, thus less attractive to herbivores (Minorsky 2019; Bell 2023). Hagihara and coworkers (2022) found that *M. pudica* leaves that had been pharmacologically and genetically manipulated to not exhibit thigmonastic rapid movements experienced twice as much herbivory by grasshoppers and caterpillars compared to unaltered plants.

**Figure 1.**
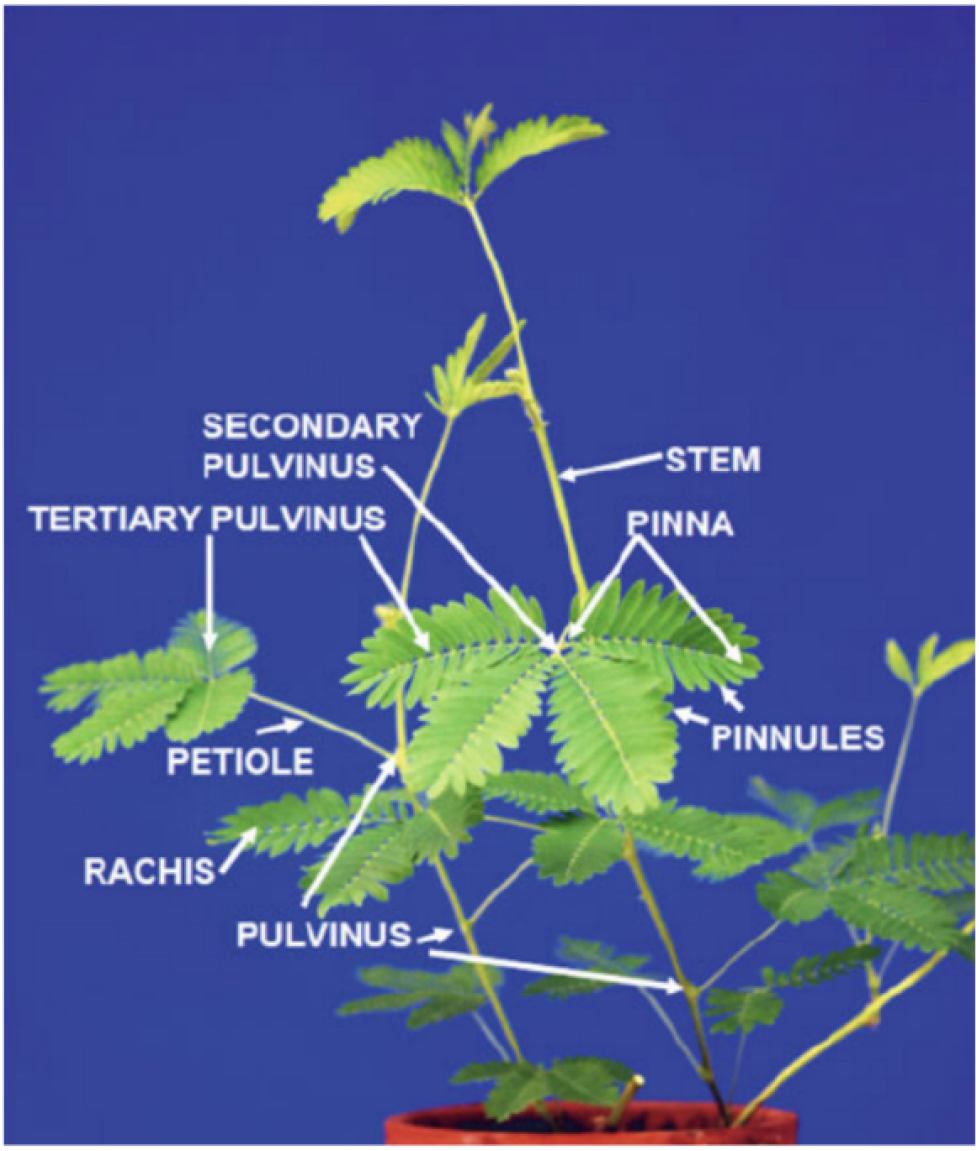
Anatomy of *Mimosa pudica* plant. Note that “PULVINUS” refers to primary pulvinus (P1). “PINNULES” are often referred to as “leaflets” (Volkov 2014).

**Figure 2.**
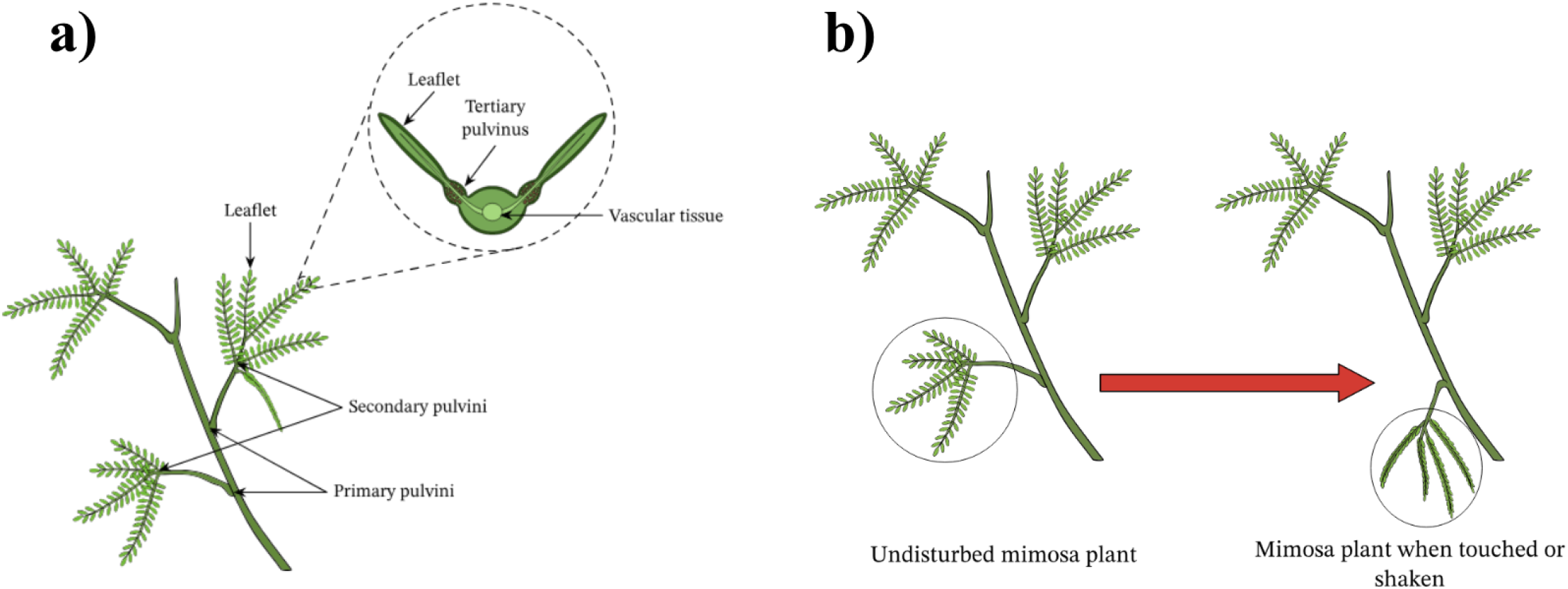
**a)** Anatomy of *Mimosa pudica* plant. **b)** Depiction of both leaflet folding (P3 activation) and drooping (P1 activation) (Nagwa n.d.).

In some studies, activations of P1-3 are collectively described as a leaf-wide “folding and drooping” response to touch stimuli; however, the thigmonastic activation exhibits more complex behavior. Hagihara and Toyota (2020) provide evidence that damaging stimuli can induce long-distance electrical signals that propagate throughout the leaf, triggering responses in areas far from the site of the initial stimulus. Thus, while P1 and P3 can activate together, this typically occurs with intense stimuli. In contrast, when non-damaging stimuli are applied to only part of the plant, P1 and P3 can activate independently. For instance, P1 can be activated by pressing on the petiole without any corresponding leaflet closure. Conversely, all the leaflets on all the pinnae of a leaf can fold inward with a touch stimulus without a P1 activation. Leaflets typically take three to ten minutes to fully reopen following P3 activation, and it takes a few minutes longer for a petiole to return to its original position after P1 activation.

Primary pulvinus activation is a reversible hydroelastic movement caused by a local change in turgor pressure, driven by ion and water redistribution (Fig. 3). Upon P1 activation, ions such as calcium move from the lower part of the pulvinus to the upper part via active transport, creating an ion concentration gradient and osmotic potential (Tran 2021; Hagihara and Toyota 2020; Volkov 2010). Consequently, water flows from the lower part to the upper part of the pulvinus, leading to a decrease in volume in the lower pulvinus and an increase in volume in the upper pulvinus. This causes the petiole to drop towards the base of the stem within two to five seconds. Following P1 activation, the ions follow the concentration gradient via a slower passive transport mechanism to reestablish ion equilibrium in the pulvinus. This causes water to slowly shift from the lower to the upper part of the pulvinus, and thus the petiole returns to its original angle in five to 15 minutes. There is evidence that a similar flux of ions and water drives the closure and opening of leaflets following P3 activation (Chen 2013; Hagihara et al. 2022).

**Figure 3.**
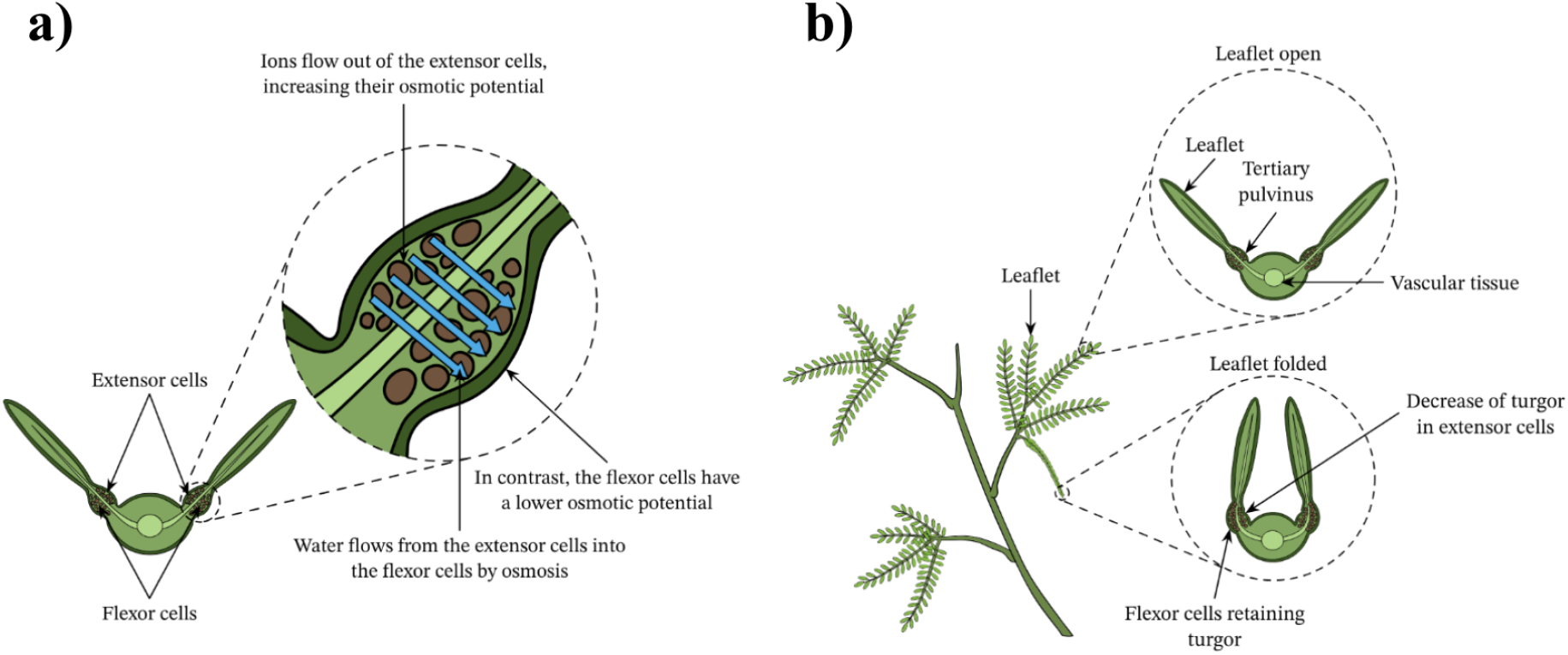
**a)** Anatomy of tertiary pulvinus. Explanation of ion/water transport during P3 activation. In P1, extensor cells = lower part of pulvinus, flexor cells = upper part of pulvinus. **b)** Depiction of P3 before and after activation as a result of a change in turgor pressure in the two parts of pulvinus (Nagwa n.d.).

The exact molecular mechanism driving the movement of calcium ions around the pulvinus is not fully understood. However, it is experimentally shown that pulvinus-mediated thigmonastic movements are energy-intensive and disrupt photosynthesis (Assmann & Zeiger 1987). Activation of the primary pulvinus (P1) is likely particularly costly energetically due to its larger size. Although the leaflets remain open, the altered leaf angle likely reduces the efficacy of photosynthesis. In contrast, activation of the smaller tertiary pulvini (P3) has a lower immediate energy cost, but results in a 40% reduction in photosynthesis rate due to the closure of the leaflets (Hoddinott 1977). Jensen (2011) found that *M. pudica* leaflets remain closed for a longer duration after P3 activation under high light conditions compared to low light conditions. This is evidence that the species balances the risks and rewards of antiherbivore behavior based on the current environmental conditions, supporting the idea that behavioral-ecology is a useful model for understanding plant responses to herbivores.

Research on primary pulvinus (P1) activation is relatively scarce - especially for repeated stimuli. My study aims to investigate P1 activation behavior in response to repeated touch stimuli in a natural setting. Existing studies have primarily focused on tracking volumetric changes within the pulvinus or the angle change following a single electrical stimulation in controlled laboratory settings (Basir 2015). I define “petiole angle change” as the total angle traveled by the petiole during P1 activation, from its starting position to its maximum change (Fig. 4b). Research on tertiary pulvinus (P3) activation has demonstrated that repeated non-damaging touch stimuli can cause the leaflets to (1) decrease their reopening time and (2) close partially rather than fully. Experimental evidence suggests that this response is a form of habituation-like plasticity rather than mechanical exhaustion (Amador-Vargas et al. 2014). Additionally, Bao and coworkers (2018) found that *M. pudica* exposed to repeated P3-activating touch stimuli for 17 days resulted in 47% lower reproductive biomass formation compared to non-stimulated plants. The tertiary pulvinus’ diminished response to non-harmful stimuli enhances the plant’s fitness by minimizing disruptions to photosynthesis and reducing rapid leaf movement energy expenditure. Given the similar hydroelastic mechanism driving both primary and tertiary pulvinus activation, it is plausible that repeated P1 activation might lead to a reduced petiole angle change after the initial stimulus. Additionally, mechanosensitive ion channels, responsible for the flux of calcium ions in pulvini, exhibit a refractory period wherein they do not carry out their function immediately after activation, even if the same stimulus is applied again (Tran 2021). It is reasonable to assume that this may manifest in a refractory period for the primary pulvinus.

**Figure 4.**
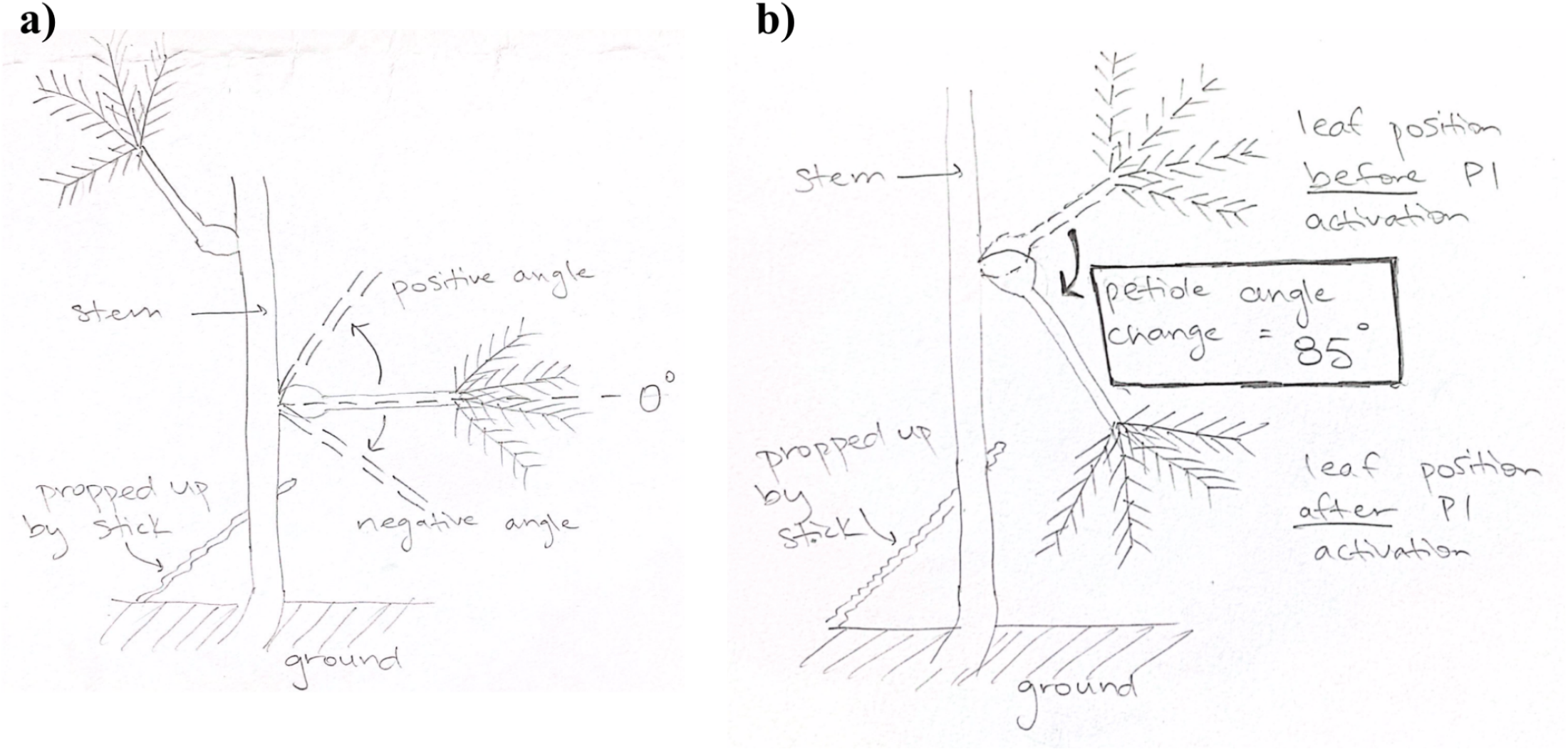
**a)** Experimental setup for both NS-P3 and P3 experiments. The stem with the leaves is propped up to vertical with a stick. This drawing can be viewed as a P.O.V. of the camera I recorded the experiments with. Note that a petiole angle of 0 degrees refers to when the petiole is perpendicular to the stem. The positive and negative petiole angles convention used in Figure 5. **b)** Experimental setup highlighting the definition of “petiole angle change” immediately following P1 activation.

Additionally, the relationship between P1 and P3 activations remains unexplored. I define “P3 activation” as the activation of all tertiary pulvini on a leaf. To my knowledge, there is no existing research on how repeated P1 activation behavior is influenced by concurrent P3 activation.

In this study, I investigate the petiole angle change with repeated P1 activating touch stimuli. I investigate whether this behavior is due to habituation-like plasticity or mechanical exhaustion. I investigate whether the addition of P3 activation affects these findings.

## MATERIALS AND METHODS

I conducted my experiments in May 2024 in Cuajiniquil, Costa Rica. The *Mimosa pudica* plants were located on a grassy soccer field on the edge of town. Since *M. pudica* behavior is influenced by light availability (Jensen 2011), all experiments were performed on plants in direct sunlight between 7-9 AM. This time frame allowed the plants to fully reopen and recover from night-time drooping before the experiments began.

On Day 1, at 6 AM, I selected stems from the same plant bunch within the same area of the soccer field. *M. pudica* is a creeping plant, so I propped up the selected stems to a roughly vertical position using sticks to better measure petiole movement without obstruction from other leaves and grasses. Only stems with more than four adult leaves were chosen, defining adult leaves as those that are dark green and have pinnae longer than two centimeters. I positioned a camera roughly one meter from the stem, perpendicular to the petioles, facing the top of the leaves to record the experiment. This setup process (Fig. 4) took approximately 15 minutes, allowing the leaves to be free from touch stimuli for 45 minutes before starting the experiment.

At 7 AM, I touched the petiole on each leaf one centimeter from the primary pulvinus with a small stick, displacing the petiole one centimeter downward towards the base of the stem. I did not touch the leaflets to avoid activating P3. I waited five minutes and then repeated the same touch stimulus. This was repeated 17 times, for a total of 18 touch stimuli per leaf. Before each stimulus, I recorded the petiole angle. P1 did not activate at every stimulus. After each stimulus, if P1 activated, I recorded the final petiole angle post-activation. Stated again, “petiole angle change” is the angle (in degrees) that the petiole traveled during P1 activation (from before activation to the end of the fast drop motion) (Fig. 4b). After the 18th stimulus, I did not touch the plant for ten minutes and recorded the petiole angle at five and ten minutes after the 18th stimulus. Following this, I applied an intense stimulus to each leaf by flicking the center of the leaf (at the base of the rachides) with my thumb and middle finger, displacing the leaf center downward by two centimeters. I will refer to this stimulus as “final flick stimulus” because it is applied after the 18th petiolar stimuli. The petiole angle was then recorded at five, ten, and 15 minutes after the flick stimulus. I refer to this set of experiments as the “NS-P3 group” (no stimulus P3 group) because I did not activate the tertiary pulvini.

On the second morning at 6 AM, the same setup (Fig. 4) was performed on the same stems/leaves of the same plant. The same stems were located using colorful string I had tied on them the previous day. The same actions were repeated as described for Day 1, with the addition that P3 was activated each time P1 activated or the leaflets were fully reopened. To activate P3, I compressed each pinna on the leaf with my thumb and index finger. I refer to this set of experiments as the “P3 group” because I activated the tertiary pulvini. On the third morning at 9 AM, the same stems were located using the colorful string. I measured their starting petiole angles and then applied the flick stimulus to each leaf, recording the petiole angle after P1 activation. I will refer to this stimulus as “initial flick stimulus” because it is applied without any preceding stimuli.

All experiments were recorded on video. Recording a petiole angle involved rewatching the video and measuring the angle with a transparent protractor on my laptop screen, approximated to the nearest five degrees. The experiments were conducted on three plant bunches/study sites. Each study site consisted of four to five stems with four to five leaves per stem, with each stem originating from a different main tap root. This amounted to 60 leaves in total (roughly 20 leaves per study site) for each of the two experimental groups (NS-P3 and P3 groups). Figure 4 depicts the experimental setup and angle measurement process. Figure 5 shows the petiole angle behavior of one leaf (for both NS-P3 and P3 groups).

**Figure 5.**
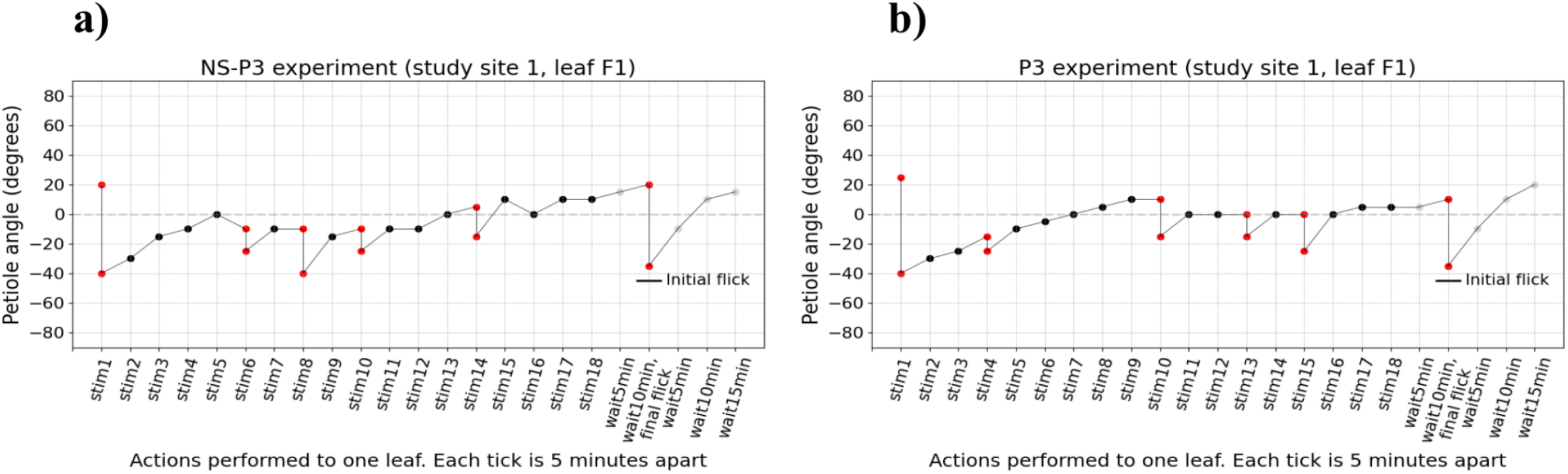
**a)** Petiole angle during NS-P3 experiment (study site 1, leaf F1). Stim1-18 refers to the 18 non-damaging stimuli applied to the petiole. Wait[5,10,15]min refers to waiting time following the last stimulus. Initial flick horizontal bar shows the petiole angle after the initial flick on Day 3 of experimentation with this leaf. Each x axis tick is five minutes apart. **b)** Petiole angle behavior for the same study site and leaf, but for the P3 experiment the following day.

Two-tailed independent sample t-tests, two-tailed paired sample t-tests, and linear regressions were performed using the python packages scipy.stats.ttest_ind, scipy.stats.ttest_rel, and scipy.stats.linregress.

## RESULTS

The highest mean petiole angle change and the highest count of P1 activations occurred at the first stimulus for both the NS-P3 and P3 groups (Fig. 6). The mean petiole angle change for all the remaining stimuli were less compared to the mean angle change at the first stimulus in both groups. The P3 group exhibited a wave-like behavior, where both the mean petiole angle change and the number of P1 activations per stimulus exhibited two troughs and two peaks.

**Figure 6.**
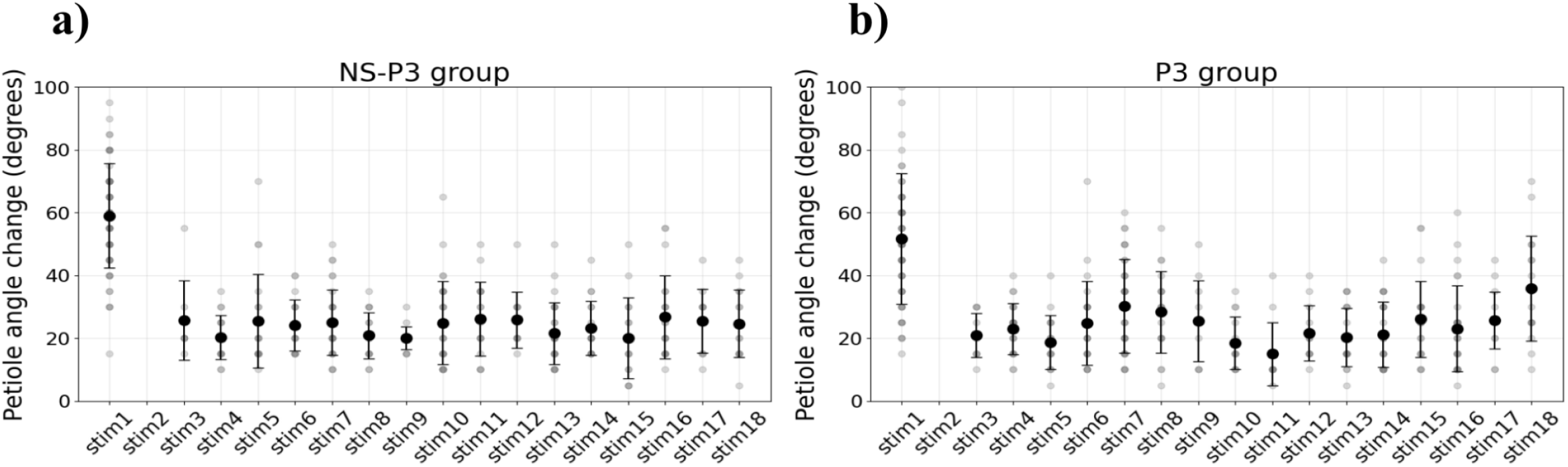
**a)** Petiole angle change for P1 activations per stimulus for NS-P3 group. Large black circles = mean of column, black bars = 1 standard deviation. **b)** Same as a) but for the P3 group.

All primary pulvini activated upon the first stimulation, resulting in a large mean petiole angle change (Fig. 6). No P1 activations occurred at the second stimulus. The mean petiole angle change at the third stimulus was lower than the first stimulus in both the NS-P3 and P3 groups (*two-tailed independent sample t-test*; p = 3.7e-6, 2.2e-5). In the P3 group, after a trough at the fifth stimulus, the mean angle change increased, reaching a peak at stimulus seven. This was followed by a trough at stimulus seven and peak at stimulus 18. Statistically significant differences were observed between the troughs and peaks (at stimuli five, seven, 11, and 18) (*two-tailed independent samples t-test*, p = 0.007, 0.003, 0.001). Although a visual inspection of this graph for the P3 group suggests a similar wave-like pattern, statistical analysis indicates that this difference is not significant.

The count of P1 activations for all leaves at each stimulus exhibited a similar oscillatory pattern in the P3 group (Fig. 7). All P1s activated at the first stimulus, and none activated at the second stimulus. For the NS-P3 and P3 groups, starting at stimulus three, the count of P1 activations increased, reaching a peak at stimulus seven. In the P3 group, the count of P1 activations exhibited a wave-like pattern with a trough at stimulus nine and a peak at stimulus 16. After stimulus seven, the NS-P3 group exhibited peaks, but exact behavior is unclear.

**Figure 7.**
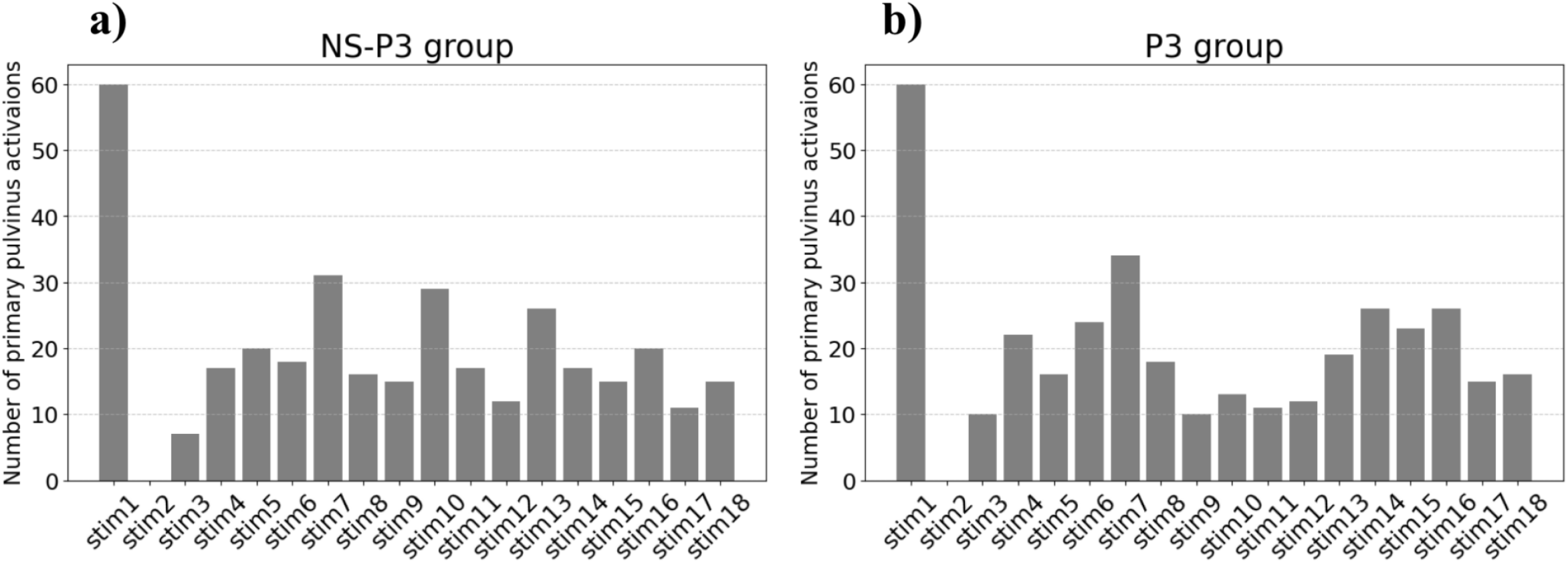
**a)** Number of P1 activations per stimulus for the NS-P3 group. **b)** Same as a) but for the P3 group.

The P1 activation behavior in the NS-P3 and P3 groups exhibited similar global behavior (Figs 6 & 7). From stimulus three to 18, the petiole angle change values oscillated around roughly the same angle for both the NS-P3 and P3 groups (*linear regression*; slopes = 0.05, 0.29; intercepts = 23.2, 20.6).

The NS-P3 and P3 groups exhibited a difference in petiole angle change at the first stimulus (Fig. 6). The mean petiole angle change at stimulus 1 was about 60 degrees for the NS-P3 group and 50 degrees for the P3 group, which is a statistically significant difference (*two-tailed paired sample t-test*; p = 0.005).

The mean petiole angle change after the flick stimulus was different between the initial flick and both the NS-P3 final flick and P3 final flick (Fig. 8). There was no significant difference between the mean petiole angle change of the final flick stimulus of the NS-P3 and P3 groups. The final flick stimulus caused P1 activation in 93% and 85% of petioles for the NS-P3 and P3 groups, respectively. The mean petiole angle change at the final flick stimulus was 29 degrees for the NS-P3 group and 25 degrees for the P3 group, which is higher than the previous P1 activation for the corresponding leaf in 87% (NS-P3 group) and 82% (P3 group) of cases. P3 activation of all leaflets was observed after the flick stimulus in nearly all leaves, indicating that the flick stimulus was more disruptive and more likely to cause P1 activation. The mean petiole angle change for the initial flick stimulus was 60 degrees, which is higher than the mean final flick stimulus for the NS-P3 and P3 groups (*two-tailed paired sample t-test*; p = 2.6e-17, 7.9e-23) (Fig. 8). The mean petiole angle change for the final flick stimulus for the NS-P3 and P3 groups have no significant difference (*two-tailed paired sample t-test*; p = 0.16).

**Figure 8.**
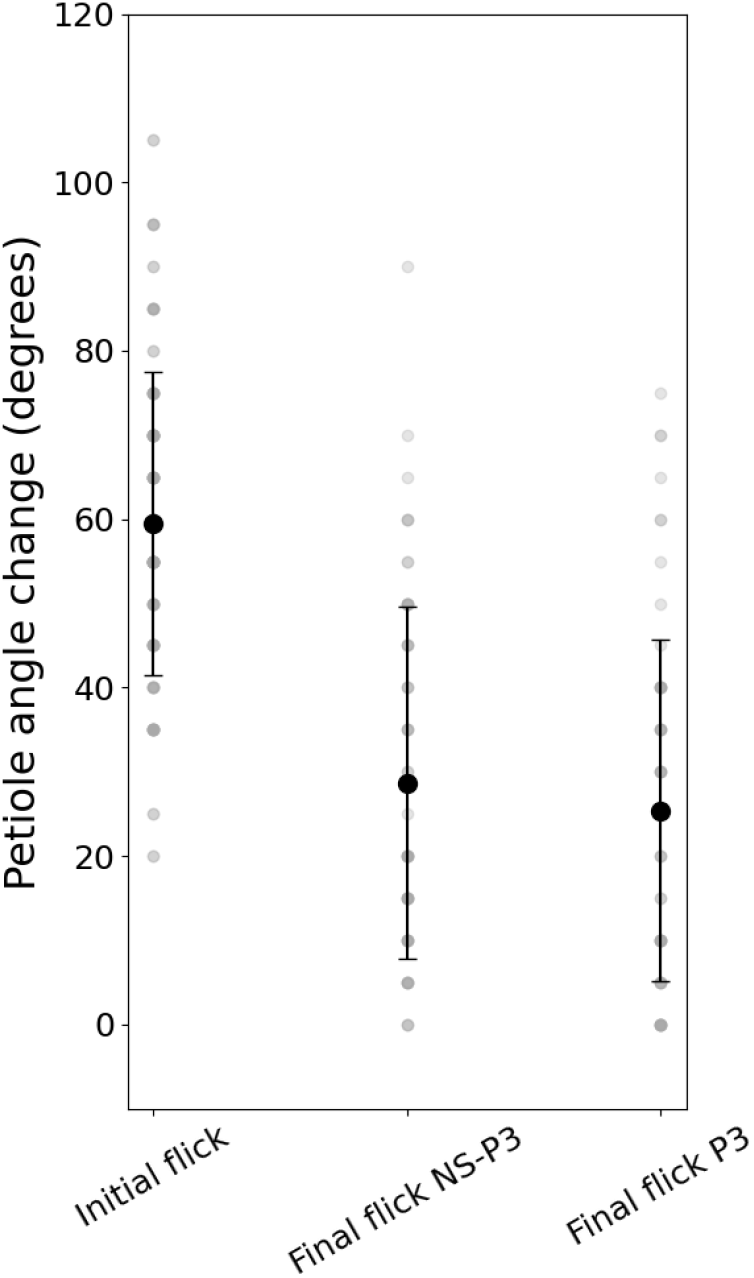
Petiole angle change after initial flick, final flick (NS-P3 group), and final flick (P3 group).

There was a range of counts of P1 activations in primary pulvini (Fig. 9). I define a low-response leaf as one that experienced 6 or fewer P1 activations during the 18 stimuli. I define a high-response leaf as one that experienced 7 or more P1 activations during the 18 stimuli. With this cutoff, 43% of leaves were low-response leaves, and 57% were high-response leaves. A single stem can have both low- and high-responsive leaves, but there is usually a dominant responsiveness type for each stem. The aforementioned results hold when looking at just low- or just high-responsive leaves.

**Figure 9.**
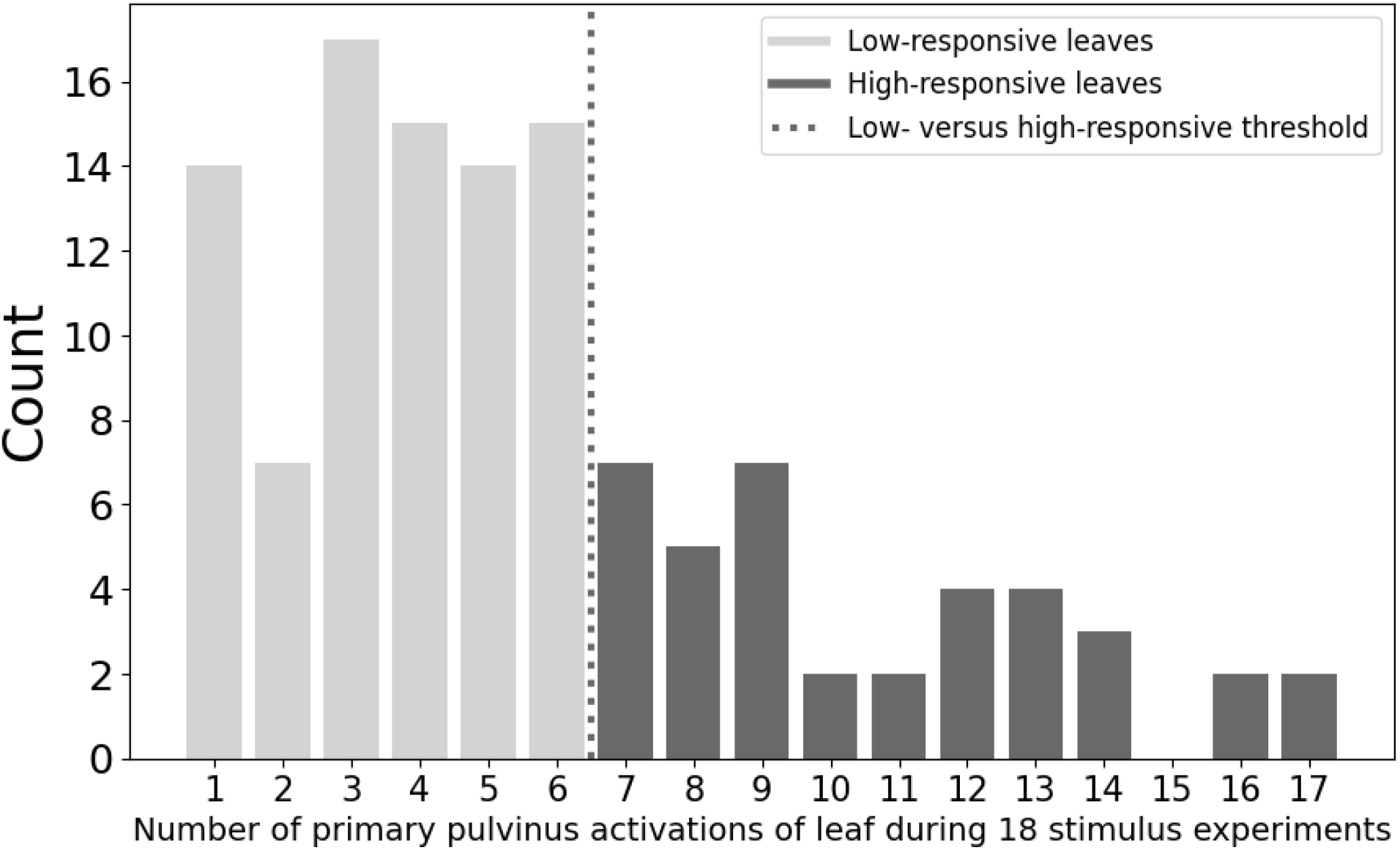
Counts of number of P1 activations across all studied leaves (both NS-P3 and P3 groups).

## DISCUSSION

### Petiole angle change and P1 activation counts per stimulus provide evidence for mechanical exhaustion in the primary pulvinus for repeated non-damaging stimuli

Both groups exhibited the highest mean petiole angle change (50 degrees for NS-P3 group and 60 degrees for P3 group) and the highest number of P1 activations at the first stimulus (Figs 6 & 7). No P1 activations occurred at the second stimulus, and the mean petiole angle change remained lower than the initial stimulus throughout the experiment for both groups. The P3 group exhibited a wave-like pattern oscillating around 20 degrees, with two troughs and two peaks, while the NS-P3 group also centered around 23 degrees but lacked statistically significant wave-like behavior.

Two main hypotheses can explain these results: (1) habituation-like response plasticity and (2) mechanical exhaustion. Habituation is the process by which an organism gradually reduces its response to repeated, non-damaging stimuli (Merchie 2023). This has been observed with full versus partial leaflet closure in adult *M. pudica* leaves (Amador-Vargas et al. 2014). In the context of thigmonastic movement driven by mechanosensitive ion channels, mechanical exhaustion is characterized by a large initial response, followed by a sustained diminished response in subsequent equivalent stimuli (Kouhen 2023). In this study, the steep decline in petiole angle change after the first stimulus, followed by a plateau at lower angle values supports the mechanical exhaustion hypothesis (Fig. 6).

The absence of P1 activations at the second stimulus further supports the mechanical exhaustion hypothesis (Figs 6 & 7). This lack of activation may result from the ions having insufficient time to return to their pre-P1 activation distribution in the pulvinus after the initial stimulus. (Tran 2021). This and the wave-like behavior in the P3 group is evidence of a refractory period in the primary pulvinus where ions are returning to their original positions, limiting the number of activations and the magnitude of angle changes after high activation periods.

Amador-Vargas and coworkers (2014) applied a more intense, damaging stimulus after the final non-damaging stimulus and saw a higher response (more full leaflet closure), using this as evidence of habituation. The same response was observed in this study where the final flick stimulus elicited a higher petiole angle change as compared to the leaf’s previous P1 activation. The flick stimulus is likely not damaging, but was shown to be more intense than the non damaging stimulus (flick was more likely to cause P1 activation, result in a higher petiole angle change during P1 activation, and trigger P3 activation in addition to P1). However, the petiole angle change for the final flick is significantly lower than the petiole angle change after the initial flick (Fig. 8). This is evidence that the primary pulvinus’ response capacity diminishes with repeated stimuli due to the energy-intensive nature of the activation and the time required for ion channels to reset, contrasting with the gradual decrease seen in habituation.

### NS-P3 and P3 groups exhibit similar overall behavior, with some differences

Both the NS-P3 and P3 groups exhibited similar overall behavior, characterized by a steep decline in petiole angle change and P1 activation counts after the first stimulus (Figs 6 & 7). However, there were notable differences in their mean petiole angle change. The P3 group’s petiole angle change was lower at the first stimulus compared to the NS-P3 group (Fig. 6). This difference could be attributed to the plant not having sufficient time to recover from the prior day’s activations, though this hypothesis is improbable given that petioles typically return to their original angles within 20 minutes. Nevertheless, without reversing the experimental order (conducting P3 on the first day and NS-P3 on the second), this hypothesis cannot be completely dismissed. The statistically significant wave-like behavior observed in the P3 group from stimuli three to 18, which was not seen in the NS-P3 group (Fig. 6), is evidence for a more pronounced refractory period in the P3 group due to the combined activation of both P1 and P3 pulvini.

### Variability in P1 activation responsiveness

The range of P1 activation counts across leaves, with some leaves being consistently low-responsive and others high-responsive, is intriguing (Fig. 9). This variation is surprising given that all studied leaves appeared similar in color and size. Generally, leaves maintained their responsiveness (low versus high) across the NS-P3 and P3 experimental days. I observed leaves of both responsiveness types on the same stem, though there was generally a dominant responsiveness type per stem. This phenomenon warrants further exploration to understand factors contributing to this differential responsiveness.

## ACKNOWLEDGEMENTS

I am immensely appreciative of Frank Joyce and Naomi Solano Guindon for their guidance in study design and manuscript preparation. I am especially grateful for their ability to inspire in me a deep wonder about the Tropics and plants.

